# A Neurophysiological Measure of Reward Sensitivity and its Limited Association with Anhedonia in Adolescents and Young Adults

**DOI:** 10.1101/467217

**Authors:** David W. Frank, Elise M. Stevens, Francesco Versace

**Affiliations:** Department of Behavioral Science, The University of Texas MD Anderson Cancer Center, Houston, Texas; The Oklahoma Tobacco Research Center, Stephenson Cancer Center, Oklahoma City, Oklahoma

**Keywords:** Anhedonia, Feedback Related Negativity, Reward, ERP, Adolescent, Young Adult

## Abstract

Anhedonia (i.e., the attenuated ability to enjoy pleasurable stimuli) characterizes multiple mood disorders, but its neurophysiological underpinnings are not yet clear. Here, we measured event-related potentials in 116 adolescents and young adults engaged in a signal detection task designed to objectively characterize the anhedonic phenotype. In line with previous studies, the behavioral results showed that approximately 35% of the sample did not develop a response bias towards the more frequently rewarded stimuli (a sign of low hedonic capacity). The event-related potentials (ERPs) evoked by the reward feedback stimuli delivered during the task showed that individuals that did not develop a response bias had significantly less cortical positivity at Fz from 224 ms to 316 ms post feedback onset compared to those that developed a response bias during the task. However, further analyses showed that this between groups difference was relatively weak, as it disappeared when we controlled for response-locked ERPs. Furthermore, the response bias observed in the signal detection task was not strongly associated with self-reported ratings of hedonic capacity. We conclude that even though the signal detection task may be used as a reward sensitivity measure in neurotypical adolescents and young adults, this task may only be able to detect clinically significant levels of anhedonia in this particular population.

## Introduction

Anhedonia, the attenuated ability to enjoy pleasurable stimuli (Chapman et al., 1976), is associated with psychiatric disorders in adults, including major depressive disorder (Loas, 1996; Treadway & Zald, 2011), bipolar disorder (Pizzagalli et al., 2008a), schizophrenia (Gard et al., 2007; Kwapil, 1998), and substance use disorder (Garfield et al., 2014; Markou et al., 1998). In adolescents, anhedonia is associated with enhanced depression severity and longer time to remission (Bennik et al., 2014; Gabbay et al., 2015; McMakin et al., 2012). Given its clinical relevance, understanding the neurophysiological mechanisms underlying anhedonia is extremely relevant as it can contribute to the improvement of diagnostic procedures and the efficacy of clinical interventions (Der-Avakian et al., 2015).

Anhedonia is generally measured using self-report questionnaires, but many of these questionnaires may not generalize to a large portion of the population. For instance, some anhedonia scales may be culturally biased and may not be relevant to all nationalities or age groups (Leventhal et al., 2006). The Snaith-Hamilton Pleasure Scale (SHAPS; Snaith et al., 1995) was specifically developed to overcome these limitations by including general statements reflecting the ability to experience pleasure during everyday events that do not depend on social class, gender, age, dietary habits, or nationality (e.g. “I would enjoy being with my family or close friends”). Due to these characteristics, the SHAPS is considered a reliable index of anhedonia in both adult (Gilbert et al., 2002; Leventhal et al., 2006) and younger populations (Leventhal et al., 2015). However, self-report measures cannot provide information about the neurophysiological underpinnings that contribute to the manifestation of anhedonia.

To supplement self-reports, Pizzagalli and colleagues (2005) developed a behavioral task aimed at objectively measuring sensitivity to reward, a key feature of anhedonia. In this signal detection task, the participant differentiates between two perceptually similar stimuli (a line drawing of a face with either a long or short mouth). During the task, a portion of correct responses are immediately followed by a reward feedback indicating a monetary gain. However, unbeknownst to the participant, one response is rewarded more frequently than the other. Due to the difficult nature of the task (the mouth lengths are nearly identical), the participant operates under high perceptual uncertainty, therefore often “guessing” the answer. In this context, individuals who are more sensitive to reward should be more likely to develop a response bias toward “guessing” the mouth length that is rewarded most often, while those who are less sensitive to rewards should be less likely to develop such a bias. Empirical findings indicate that individuals with high scores on self-report scales of anhedonia are less likely to develop a response bias towards the frequently rewarded stimulus compared to controls (Bogdan & Pizzagalli, 2006; Pizzagalli et al., 2008a; Pizzagalli et al., 2008b; Pizzagalli et al., 2005; Santesso et al., 2008a).

One advantage of this signal detection task is that it allows researchers to record neurophysiological responses to the stimuli indicating monetary gains, specifically the amplitude of the feedback related negativity (FRN; Cohen et al., 2007). Typically, the FRN is described as a negative-going component of the event-related potential (ERP) that peaks between 200 and 400ms post-stimulus onset. This ERP component has been suggested to reflect performance monitoring, as it is generated when performance expectations are violated by feedback indicating unexpected outcomes (Hajcak et al., 2005; Holroyd & Coles, 2002; Holroyd & Coles, 2008; Oliveira et al., 2007). The results that Santesso et al. (2008a) obtained recording ERPs during the signal detection task described above seem in line with this interpretation. Santesso et al. (2008a) showed that the feedback stimuli signaling monetary gains evoke a larger FRN in individuals who do not develop a response bias (i.e. are less sensitive to rewards) relative to individuals who develop a response bias (i.e., are sensitive to rewards). The amplitude of the FRN also correlates with higher levels of anhedonia (Santesso et al., 2008a), and is enhanced for individuals with major depressive disorder compared to controls (Mueller et al., 2015; Santesso et al., 2008b). Taken together, these electrophysiological and behavioral results are promising because they provide objective measures of anhedonia that are not possible when solely relying on self-report measures. Additionally, measuring the FRN offers the opportunity to investigate the neural mechanisms that underlie anhedonic symptomology. Thus, combining information from electrophysiological, behavioral, and self-report measures can provide a more accurate assessment and, ultimately, improve treatments for individuals with anhedonia.

While these studies provide insight into reward learning, they were conducted with college aged adults and it is unclear the extent to which they translate to younger individuals. Due to the association between anhedonia and both increased depression severity and longer time to remission in the adolescent population (Bennik et al., 2014; Gabbay et al., 2015; McMakin et al., 2012), it is critical that the assessment of anhedonia in adolescents is as accurate as possible. However, electrophysiological research in this population is limited. We thus sought to replicate the neurobehavioral results from Santesso and colleagues (2008a) while also extending our focus towards younger individuals.

Here we collected self-report, behavioral, and electrophysiological data to assess the multiple components contributing to anhedonia in adolescents and young adults. After completing questionnaires, participants performed the signal detection task while we continuously collected electroencephalogram (EEG). We hypothesized that individuals with reduced reward sensitivity, (as measured by their performance on the signal detection task) would report greater anhedonia scores on the SHAPS and exhibit a less negative FRN to reward feedback stimuli delivered during the signal detection task.

## Methods

### Participants

All study procedures were approved by the Institutional Review Board at the University of Oklahoma Health Sciences Center. A total of 122 participants were recruited at the University of Oklahoma Health Sciences Center’s Oklahoma Tobacco Research Center. Due to technical problems, data from six participants were not usable, hence 116 subjects were included in the final data set. Participants were young individuals (14-21 years old, 17.2 avg.; 60 female/55 male) recruited using flyers distributed at local community centers and by contacting local high schools. Participants reported being in good health and free of any psychological disorders. Written consent was obtained from all participants. For participants under the age of 18 (n=61), we also obtained in-person consent from a legal guardian. In addition to the procedures described below, the participants also viewed a slideshow depicting various naturalistic scenes. Those data are being analyzed and will be published elsewhere.

### Questionnaires

Before participating in the EEG session, participants completed the following questionnaires: the Barratt Impulsivity Scale (BIS; Patton et al., 1995), Positive and Negative Affect Scale (PANAS; Watson et al., 1988), and the Snaith-Hamilton Pleasure Schedule (SHAPS; Snaith et al., 1995). The BIS is designed to measure impulsiveness by including eight items related to impulsive or non-impulsive behaviors and preferences (e.g. “I concentrate easily”). The PANAS measures positive and negative affect where participants endorse a series of 20 items (e.g. “During the past week I felt upset”). The SHAPS was specifically designed to measure anhedonia across age and cultural groups and contains 16 items for participants to endorse (e.g., “I would get pleasure from helping others”). The SHAPS was coded from zero to three, with three denoting the highest level of anhedonia on each item and zero denoting no anhedonia. Participant gender, age, and questionnaire scores are reported in Table 1. To assess the association between reward sensitivity and anhedonia, for each questionnaire we conducted a one-way ANOVA between learners and non-learners (see Reward bias and task performance, below) using the average score for each participant.

**Table 1.**
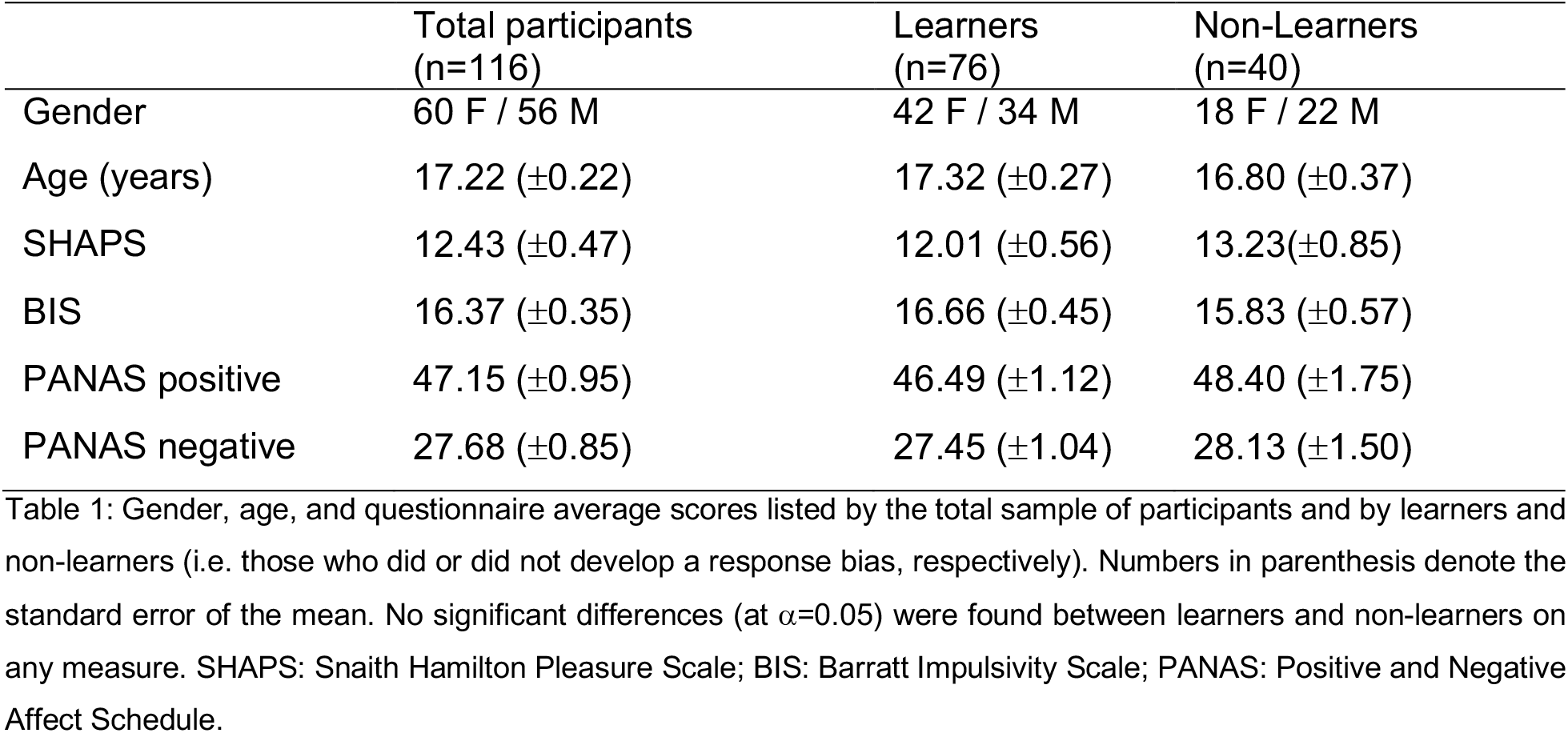
Participant gender, age, and questionnaire scores.

listed by the total sample of participants and by learners and non-learners (i.e.
|  | Total participants<br>(n=116) | Learners<br>(n=76) | Non-Learners<br>(n=40) |
| --- | --- | --- | --- |
| Gender | 60 F / 56 M | 42 F / 34 M | 18 F / 22 M |
| Age (years) | 17.22 ($\pm 0.22$) | 17.32 ($\pm 0.27$) | 16.80 ($\pm 0.37$) |
| SHAPS | 12.43 ($\pm 0.47$) | 12.01 ($\pm 0.56$) | 13.23 ($\pm 0.85$) |
| BIS | 16.37 ($\pm 0.35$) | 16.66 ($\pm 0.45$) | 15.83 ($\pm 0.57$) |
| PANAS positive | 47.15 ($\pm 0.95$) | 46.49 ($\pm 1.12$) | 48.40 ($\pm 1.75$) |
| PANAS negative | 27.68 ($\pm 0.85$) | 27.45 ($\pm 1.04$) | 28.13 ($\pm 1.50$) |

### Signal detection task

Before starting, a research assistant described the experiment to the participant, and explained that the task offered the possibility to gain real money that would be paid at the end of the session. Eight practice trials were then completed (with the option to repeat the practice trials) to make sure that the participants understood the procedure. The signal detection task was implemented using E-prime software (version 2.0; Psychology Tools, Pittsburgh, PA) and the stimuli were presented on a LCD screen placed approximately 60 cm from the participant.

In each trial (Fig. 1), a line-drawn face with no mouth would appear on the screen for 500ms, followed by a long or short mouth superimposed for 100ms. The participant then had up to 1500ms to decide if they had seen a short or a long mouth by pressing the “f” or “j” key on the keyboard (key assignment was counterbalanced across participants). The key press was followed by a blank screen on 60% of trials, while on 40% of trials the key press was followed by feedback indicating that the participant had won five cents. Unbeknownst to the participant, one of the two answers (either the long or the short mouth, counterbalanced across participants) was rewarded three times more frequently than the other (90 out of 120 total feedback trials). To ensure that the 3:1 proportion of trials in the “rich” (i.e., the condition rewarded more often) and “lean” (the condition rewarded less often) conditions would remain the same for all participants, if the participant chose the wrong response in a trial designated to be followed by correct feedback, that trial was repeated and feedback was withheld until the participant answered correctly. The task included three blocks with a short break between blocks to allow the participant to relax. Each block included 40 trials followed by the feedback, randomly interspersed among 60 trials followed by the blank screen (the total number of trials within each block was 100, plus any repeated incorrect trial), and each trial was followed by a variable interval between 500 and 1500 ms.

**Figure 1:**
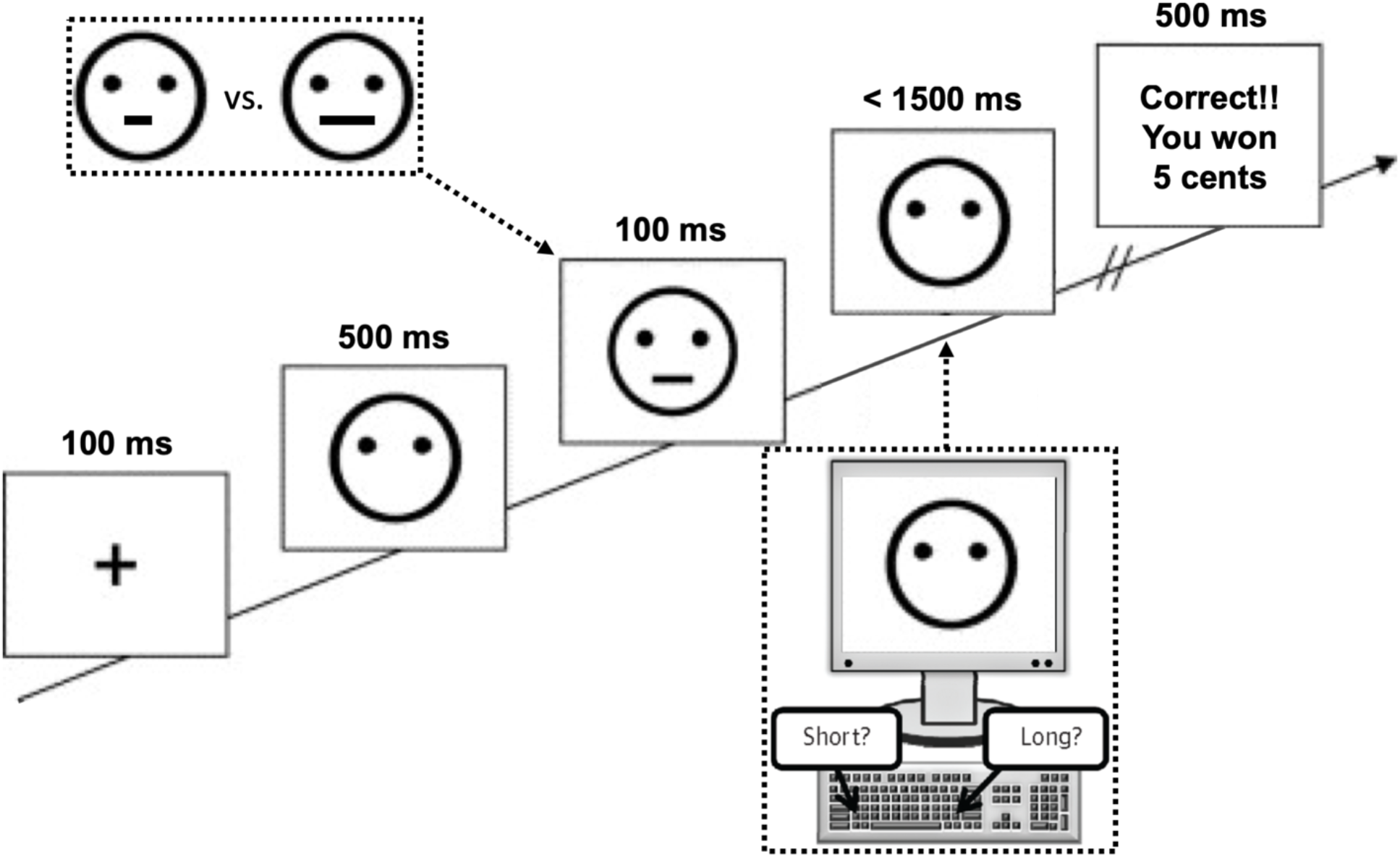
Signal detection task. In each trial, a line-drawn face missing the mouth would appear on the screen for 500ms, followed by a long or short mouth presented for 100ms. The participant then had up to 1500ms to decide if they had seen a short or long mouth by pressing the “f” or “j” key on the keyboard. The key press was followed by a blank screen on 60% of trials (not shown here), while on 40% of trials the correct key press was followed by a feedback (shown here) indicating that the participant had won 5 cents. Unbeknownst to the participant, one of the two correct answers (either the long or the short mouth, counterbalanced across participants) was rewarded 3 times more frequently than the other (90 out of 120, or 75%, of total feedback trials). Note that difference in mouth lengths depicted in the top left of the figure are embellished for illustration purposes; the actual mouth lengths during the task were nearly perceptually identical. Adapted from Pizzagali et al., 2005.

### Reward bias and task performance

In order to determine individual performance and identify individuals that developed a bias towards the more frequently rewarded stimulus (i.e. either the long or the short mouth), for each block we calculated discrimination and bias index scores applying the formulas reported in Figure 2 (Panels A and B). We then subtracted the bias index score of block one from the bias index score of block three. As a result, scores greater than zero denote individuals that, during the task, develop a progressively stronger response bias toward the condition reinforced more often. We labeled these individuals as learners. Scores less than or equal to zero indicate individuals that do not develop a bias toward the more frequently reinforced condition. We labeled these individuals as non-learners. We also computed the discrimination index for each block in order to ensure that bias changes were not the consequence of changes in performance accuracy. Since all participants received the same number of feedback trial with the preset 3:1 ratio, to avoid biasing the results, all computations to calculate discrimination and bias excluded trials followed by a feedback.

**Figure 2:**
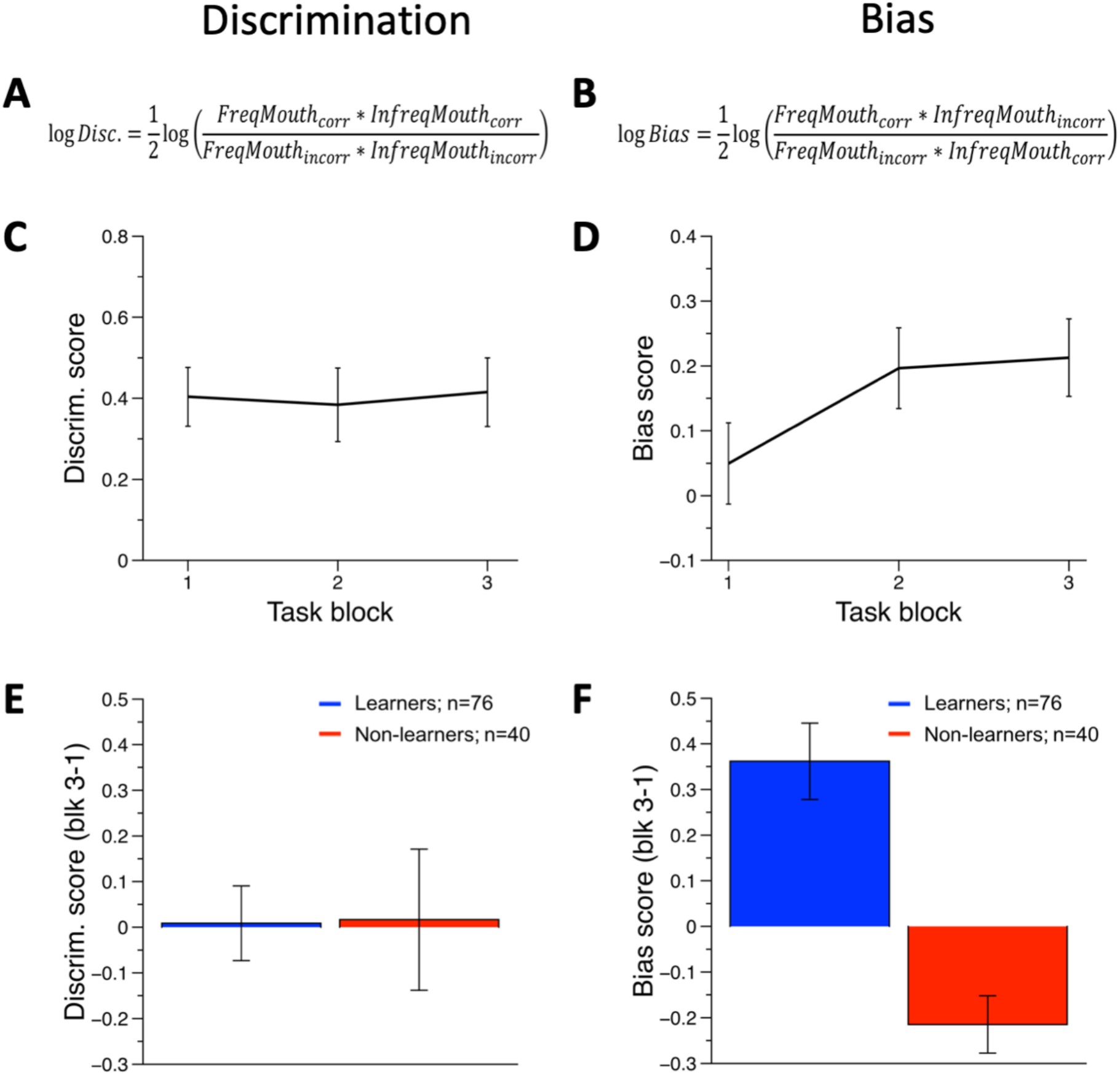
On average, using discrimination and bias formulas (A & B, respectively), participants showed no improvement in performance across the duration of the task (C) and a tendency to develop a bias towards more towards frequently rewarded response (D). However, 40 participants did not develop a bias towards the frequently rewarded response. Participants were labeled as learners if they scored greater than zero when the bias values of block one were subtracted from block three, while those who scored less than or equal to zero were labeled as non-learners. These difference scores are plotted for both discrimination (E) and bias (F). Discrimination (Disc.) was calculated using the formula shown in panel A to determine task performance, with a score of zero denoting random stimulus selection. Bias was calculated using the formula shown in panel B to determine the tendency to select the more frequently rewarded stimulus over the less frequently related stimulus, with a score of zero denoting no bias. Calculations included the number of correct (corr) and incorrect (incorr) responses to frequent (freq) and infrequent (infreq) mouth lengths that were not followed by feedback stimuli. Note that every participant received the same number of feedback trials (120) with a fixed ratio of 3:1 between the rich and lean conditions. These trials were excluded from the analyses to avoid distorting the results. Error bars denote 95% confidence intervals.

### EEG acquisition and data reduction

We recorded continuous EEG using a 32-channel actiCAP system (Brain Products GmbH, Gilching, Germany) and sampled at 500 Hz with electrodes positioned according to the 10/20 system. We then referenced each pre-amplified electrode to Cz. We set software filters to a 100Hz high cutoff and a 60Hz notch, and re-referenced the data offline to the average reference and corrected eyeblinks using Brain Electrical Source Analysis software (BESA GmbH, Gräfelfing, Germany). We then exported the data to BrainVision Analyzer (Brain Products GmbH, Gilching, Germany), where the data were filtered using 40 Hz low-pass and 0.1 Hz high-pass cutoffs, segmented (300ms pre- to 800ms post-response onset), and baseline corrected from 200ms prior to the response. Because, in this task, the feedback appears immediately after the response, time locking the ERPs to the response is effectively the same as time locking the ERPs to the feedback onset. We marked each segment as contaminated by artifacts if the waveform had a voltage step greater than 10 µV/ms, a maximal voltage difference greater than 70 µV over 50ms, less than 0.5 µV over 100ms, or amplitudes above 100 µV or below −100 µV. We interpolated channels using their four nearest neighbors if greater than 40% of a channel’s segments were contaminated by such artifacts.

### ERP Analysis

We focused our analyses on the ERPs observed at channel Fz. We selected this channel because previous work (Santesso et al., 2008a) indicates that Fz is the site where the difference in the amplitude of the ERPs evoked by feedback stimuli signaling rewards recorded in learners and non-learners is the largest. Furthermore, following Santesso et al., we computed the ERPs including only trials in the rich condition (i.e., the most frequently reinforced condition). To avoid biasing the ERP estimates measuring peak differences within arbitrary time windows (Clayson et al., 2013; Keil et al., 2014; Luck, 2005a; Luck & Gaspelin, 2017), to assess ERP differences between learners and non-learners, we computed a one-way ANOVA at each time point from 300ms to 800 ms post feedback onset.

As mentioned above, during the signal detection task, the feedback appears immediately after the response. Hence, the electrocortical responses evoked by the feedback may be confounded with those evoked by the motor response. To isolate the electrocortical activity specifically evoked by the feedback stimuli, we took advantage of the fact that, during the task, 60% of the responses are not followed by a feedback stimulus. Hence, for each participant, we calculated the ERPs evoked by the trials not followed by feedback stimuli and subtracted them from the ERPs evoked by the trials followed by the feedback stimuli. This resulted in a difference waveform corrected for motor response for every participant. To test for the presence of differences between learners and non-learners on these difference waves, we computed a oneway ANOVA at each time point from 300ms to 800 ms.

### Age-related analysis

To determine how age influenced results, we divided participants into terciles. We calculated terciles to maximize the difference between age groups while maintaining a suitable number of participants with equal group sizes. The oldest tercile group consisted of 19, 20, and 21-year old participants. The youngest tercile group consisted of 14 and 15-year old participants. Each of these groups included 37 participants.

## Results

### Behavioral paradigm

We first evaluated participant task performance across blocks. As seen in Figure 2c, participants’ discrimination of the correct stimulus did not improve across the three blocks. As mentioned above, it was critical that participants performed this way because a bias towards the more frequently reinforced stimulus can only develop when the participant is uncertain of the correct answer. In fact, on average, participants *did* develop a bias toward the more frequently rewarded stimulus (Fig. 2d). Furthermore, as expected, this trend was not present in all of the participants; after subtracting the bias scores of block one from block three, a total of 76 individuals showed a positive difference (learners) and 40 individuals showed a negative difference (non-learners; Fig. 2f). The discrimination score remained constant across the blocks and comparable in both groups (Fig. 2e).

### Questionnaires

We next tested the relationship between reward learning and self-reported anhedonia. While non learners had slightly higher anhedonia scores than learners on the SHAPS, this difference was not significant for neither SHAPS scored from zero to three (Fig. 3; F_(1,114)_=1.46, p=0.23), nor from zero to one (F_(1,114)_=0.50, p=0.48). This might be due to our sample consisting of healthy individuals with a relatively narrow range of self-reported anhedonia scores. We also found no group differences in scores on the BIS (F_(1,114)_=1.34, p=0.25), PANAS positive (F_(1,114)_=0.63, p=0.43), or PANAS negative (F_(1,114)_=0.12, p=0.73). Therefore, in this particular pool of healthy participants, it appears that reward learning does not differentiate individuals with high versus low anhedonia, as measured by the SHAPS.

**Figure 3:**
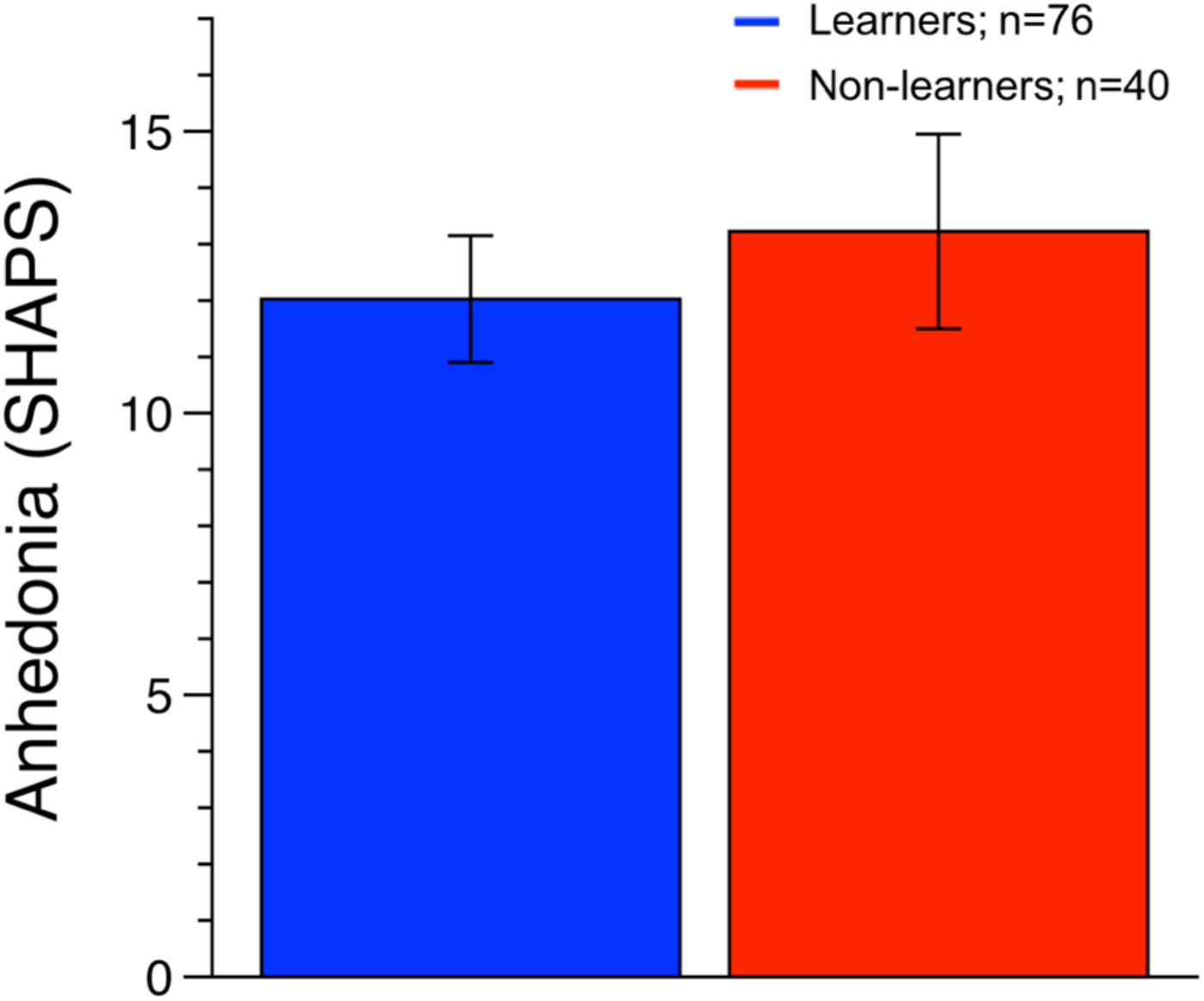
Learners and non-learners show no statistically significant difference on the SHAPS assessment of anhedonia. Learners who developed a response bias towards the more frequently rewarded stimuli show no statistically significant distinction on the Snaith-Hamilton Pleasure Scale from non-learners who did not develop a bias (F_(1,114)_=1.46, p=0.23). This might be due to the narrow range of anhedonia scores in our healthy sample. Error bars denote 95% confidence intervals.

### ERPs

In order to assess electrocortical differences between learners and non-learners, we conducted at channel Fz a point by point analysis from 300 ms before feedback onset to 800 ms post-feedback onset. As shown in Figure 4a, the feedback stimuli prompted less positivity in the ERPs of non-learners than learners and this difference crossed the (*uncorrected for multiple comparisons*) statistical threshold of p<.05 at 224 ms and remained above this threshold until 316 ms.

**Figure 4:**
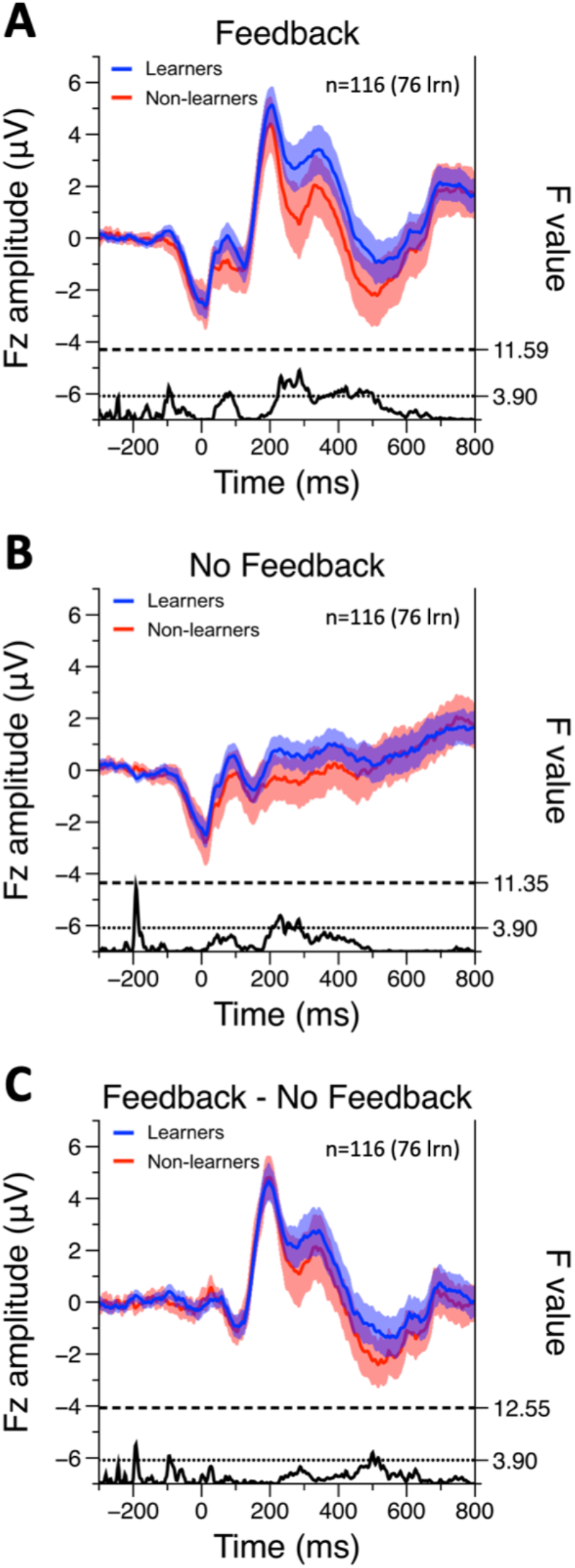
Non-learners show an enhanced Feedback Related Negativity only when including the volitional response to feedback. Channel Fz grand averaged waveforms are plotted for learners and non-learners after they received the more frequently rewarded (rich) feedback (A), no feedback (B), and no feedback trials subtracted from feedback trials (C). The solid black line at the bottom of each panel represents the pairwise one-way ANOVA conducted at each time point. ERP amplitudes and F-values of the point by point analysis are labeled on the left and right y-axes, respectively. The p=.05 threshold (F=3.9), uncorrected for multiple comparisons, is denoted by the dotted horizontal black line. The dashed horizontal black line denotes the p=.05 threshold corrected using randomization tests (10,000 random permutations run on all data points in the time series); this was not used during statistical analysis, but is provided for reference. Shaded areas denote 95% confidence intervals.

### Age-related and gender effects

To determine the extent to which age influences the results, we divided the whole sample into terciles and compared the oldest (19, 20, and 21 years old) and youngest (14 and 15 years old) terciles. The proportion of learners to non-learners was not statistically different between the upper and the lower tercile (X^2^=0.668, p=0.41; Table 2). We found that older participants endorsed lower scores of anhedonia on the SHAPS than the younger participants (F_(1,72)_=14.20, p<0.001). The ERP analyses conducted on each age group replicated the results observed in the whole sample: while feedback stimuli evoked slightly lower FRNs (i.e. more positivity) in learners than non-learners, this difference never approached significance. Furthermore, both males and females had similar bias scores (F_(1,112)_=0.004; p=0.95), and the gender by learner/non-learner interaction was not statistically significant (F_(1,112)_=1.96, p=0.16).

**Table 2.**
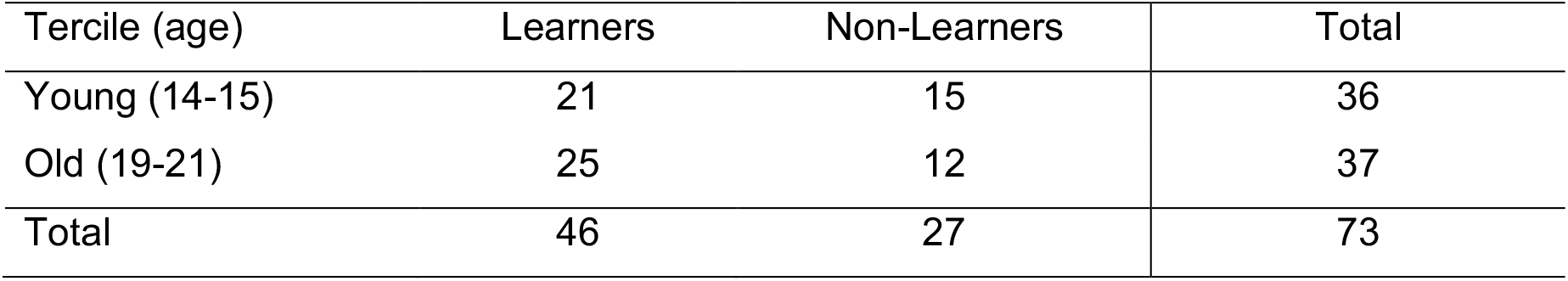
Number of participants in the oldest and youngest tercile groups.

| Tercile (age) | Learners | Non-Learners | Total |
| --- | --- | --- | --- |
| Young (14-15) | 21 | 15 | 36 |
| Old (19-21) | 25 | 12 | 37 |
| Total | 46 | 27 | 73 |

## Discussion

Here we sought to replicate previous work supporting the use of ERPs evoked by reward feedback stimuli during a signal detection task as an electrocortical index of anhedonia. We successfully used the signal detection task to identify young participants who were either sensitive or insensitive to reward. However, unlike previous studies (Santesso and colleagues (2008a), those who were relatively insensitive to reward did not report significantly lower anhedonia scores compared to their reward sensitive peers. We successfully replicated previous results showing that reward feedback evoked less positive ERPs in non-learners relative to learners, however this difference was only marginally significant. Furthermore, when we did take into account the influence that motor responses had on the ERPs, the voltage difference between learners and non-learners almost disappeared and did not approach statistical significance (Fig. 4c).

### ERPs

We implemented the signal detection task following established procedures (Pizzagalli et al., 2005; Santesso et al., 2008a) and the feedback stimuli were presented as soon as the participant pushed the response button. As a consequence, the ERPs evoked by the feedback stimuli are confounded with the ERPs evoked by the responses. To remove the contribution of the motor response from the ERPs generated by the feedback stimuli, we computed difference waves (reference to probably a Steve luck article or book chapter where it explains the idea of computing difference waves). By isolating the ERP components present exclusively after feedback presentation, we expected to magnify the electrocortical differences between learners and non-learners. However, upon conducting this subtraction, the electrocortical differences between learners and non-learners no longer approached statistical significance, even when no correction for multiple comparisons was applied. While this outcome was unexpected, similar findings have been presented by Weinberg and colleagues (2012). These researchers implemented a task in which participants had to make a decision and received feedback either one second after their motor response or six seconds after their motor response. They found that the FRN was present after one second, but eliminated after six seconds. The authors suggested that the elimination of the FRN after an extended delay may be due to interference between maintaining response-feedback contingencies. Furthermore, as our results indicate, the individuals that develop a response bias have ERPs that are more positive than in individuals not developing a response bias even, when no feedback is provided (see Figure 4C). Hence, the FRN differences observed when feedback stimuli are delivered immediately after the response seem to depend on a contribution of both feedback and response.

Although beyond the scope of the current work, we should also mention that in the past few years researchers proposed a different interpretation of ERPs evoked by feedback stimuli. Initially, the FRN was described as a negative-going component that followed feedback stimuli indicating negative outcomes such as errors or losses (Luck, 2005b; Miltner et al., 1997). However, the results from more recent studies focusing on how gains affect the FRN (Holroyd et al., 2008; Proudfit, 2015) suggest that rewards generate a positive-going component (called the reward positivity, or RewP), rather than losses generating a negative-going component (Foti et al., 2011; Liu et al., 2014). The authors reached this conclusion by considering another component that is generated by task-relevant stimuli (i.e., both gain and loss feedback stimuli): the N200. The N200 is a negative-going component that has a similar temporal evolution and spatial distribution as the FRN (Baker & Holroyd, 2008; Baker & Holroyd, 2011; Gehring & Willoughby, 2002). Applying PCA analyses on the ERPs evoked by both losses and gains, Foti and colleagues (2011) demonstrated that while both gains and losses evoke an N200, only gains evoke the RewP. However, due to temporal and spatial overlap, the RewP is masked by the N200. The characteristics of the signal detection task used here, where participants only received feedback stimuli signaling positive outcomes, seems more compatible with this interpretation. Our results should be interpreted as showing that the feedback stimuli predicting rewards evoke more positive ERPs in people that develop a bias (i.e., the people more sensitive to rewards) than in people that do not develop the bias. This hypothesis could be tested in future studies by introducing negative feedback trials to the signal detection task. If the results observed here are due to larger RewP evoked by individuals sensitive to rewards, then the FRN (or, rather, the N200) to the negative outcomes should be similar in both groups.

### Pleasure scale

Contrary to our initial hypothesis, we did not find differences in reward sensitivity, as measured by the signal detection task, between individuals who endorsed high versus low anhedonia scores on the SHAPS. This was surprising because enhanced FRN amplitude (i.e. more negativity) has been found both in individuals with depression (Foti et al., 2014; Liu et al., 2014) and in healthy individuals endorsing higher depression scores than their peers (Foti & Hajcak, 2009). However this is not always the case, as work by Chen and colleagues (2018) recently found no differences in FRN amplitude between anhedonic (determined using the Temporal Experience of Pleasure Scale; Gard et al., 2006) and healthy control participants for either reward or punishment contexts. It is also possible that, for healthy adolescents, the SHAPS questionnaire may not be sensitive enough to correlate with the subtle differences in reward sensitivity evinced by the signal detection task.

## Conclusion

In sum, given our relatively large sample size, we were able to closely replicate the results found by Santesso and colleagues (2008a) in adults; individuals that did not develop a bias during the signal detection task had less positive ERPs within a time window compatible with the FRN, compared to people who did develop a bias. However, these ERP differences were relatively weak and only met the conventional p<0.05 threshold when no correction for multiple comparisons was applied. This is underscored by the fact that when we removed the confound of the motor responses in order to exclusively examine the effects due to feedback, the ERP negativity difference between groups essentially disappeared. This was surprising, as we expected *more* reliable differences to emerge from the ERPs between groups when both removing potential confounds and including a large sample size.

The rates of major depressive disorder are high among adolescents (Hankin et al., 1998; Oldehinkel et al., 1999) and anhedonia is a distinct predictor of this disorder (Bennik et al., 2014; Gabbay et al., 2015; McMakin et al., 2012), which also predicts higher rates of substance use among the adolescent population (Luby et al., 2018). It is therefore of critical importance to understand the neurophysiological underpinnings of anhedonia and to continue to develop markers of depressive mood disorders. Although the difference between groups were weak, it is still possible that the signal detection task may be successfully implemented in young individuals to objectively assess individual differences in hedonic capacity. As mentioned above, our study only included healthy adolescents, and the signal detection task may yield better results when comparing healthy controls to individuals with a clinically significant depressive mood disorder. This task may be additionally useful as it is an objective measure of reward sensitivity and may therefore be used to measure a distinct endophenotype associated with depression (Hasler et al., 2004). Thus, future studies investigating anhedonia among adolescent populations should continue to explore the utility of the signal detection task while incorporating electrophysiological data collection to capture the non-conscious elements of anhedonia.

## Notes

Funding information: This work was supported by the National Institute on Drug Abuse under awards R01-DA032581 and R21-DA038001 to Francesco Versace and by the National Institute of General Medical Sciences under award U54GM104938 to the Oklahoma Shared Clinical and Translational Resources.

